# pileup.js: a JavaScript library for interactive and in-browser visualization of genomic data

**DOI:** 10.1101/036962

**Authors:** Dan Vanderkam, B. Arman Aksoy, Isaac Hodes, Jaclyn Perrone, Jeff Hammerbacher

## Abstract

pileup.js is a new browser-based genome viewer. It is designed to facilitate the investigation of evidence for genomic variants within larger web applications. It takes advantage of recent developments in the JavaScript ecosystem to provide a modular, reliable and easily embedded library.

**Availability:** The code and documentation for pileup.js is publicly available at https://github.com/hammerlab/pileup.js under the Apache 2.0 license.

## 1. Introduction

As sequencing has become increasingly ubiquitous, there has been a proliferation of variant calling programs. Before a physician or researcher takes action on a variant, however, it remains essential to inspect the evidence for it manually. Track viewer visualizations such as those provided by UCSC, IGV and Dalliance have long been the preferred way to do this (Kent et al., 2002; Robinson et al., 2011; Down et al., 2011). As data and workflow management systems move to the browser, it becomes increasingly appealing to embed these visualizations directly within larger web applications. This results in reduced latency, allows extensive customization and avoids the cognitive overhead of context switching between applications.

Here, we describe pileup.js, a JavaScript library for interactive and in-browser visualization of genomic data. pileup.js is extensively tested, uses a modern code base that is oriented towards re-usability and performance, and is well-documented for easier customization and usability by other developers. The latest version, v0.6.1, supports visualization of genomic tracks for reference sequences, mapped reads (paired or unpaired), read depth, variants and gene annotations. pileup.js was initially developed to be embedded within the Cycledash variant inspector, but it can be used within any web application. (Hodes et al, 2016)

## 2. Methods and Technologies

Driven by the ubiquity of web browsers, the JavaScript development ecosystem has seen a maelstrom of activity over the past several years. This has resulted in major new technologies and radically different approaches to solving problems. Rather than adapting existing systems to these new tools, we elected to create pileup.js, which is built from the ground up for today’s JavaScript ecosystem. We highlight a few of the most important tools here:

- ES2015 is the latest version of the ECMAScript (JavaScript) language (E. ECMAScript, 2015). It solves many long-standing issues, e.g. the difficulty of defining class hierarchies and the lack of a module system. We embrace these features and use babel and browserify to make them work in current-generation web browsers.
- We pull in third-party dependencies using the Node Package Manager (NPM). This allows us to take advantage of “battle hardened” code written by other developers for tasks that are not specific to pileup.js, e.g. inflating gzipped data.
- We use React.js for managing state within the genome viewer. This ensures that state changes (e.g. panning, zooming and toggling options) are consistently reflected throughout the user interface.
- We use the FlowType static analysis system to ensure the type safety of our code (Chaudhuri et al, 2014). Static type systems clarify the inputs and outputs of functions, catch errors and facilitate refactoring. Users of pileup.js can choose to use its type definitions as they wish.

In addition to using contemporary web technologies, our development process is designed to lead to higher-quality software. All code is unit-tested and goes through a blocking review by a peer. This ensures that previously-fixed bugs do not regress and that all code was understandable to at least one developer who did not write it. This leads to better documentation and cleaner APIs for pileup.js.

pileup.js uses the HTML5 canvas to render its visualizations. It uses off-screen buffers to achieve faster drawing and smoother panning. We chose canvas over other technologies (e.g. SVG) because it led to simpler code and better performance. We initially used SVG but were able to gain a 5x performance improvement by migrating to canvas (Vanderkam, 2015).

Like dalliance and IGV, pileup.js loads data over HTTP using Range requests, which are widely supported by popular servers such as Apache and nginx. For some data sources, e.g. BAM files, it may require several serialized requests to load all the information for a single genomic region (Li et al., 2009). In this case it is more efficient to situate the data loading logic closer to the data itself, to reduce the round trip time. This can be achieved by running a GA4GH server. pileup.js supports v0.5 of the GA4GH protocol (Terry et al., 2014). Support for other data loading schemes can be added via user-defined sources.

**Figure 1.**
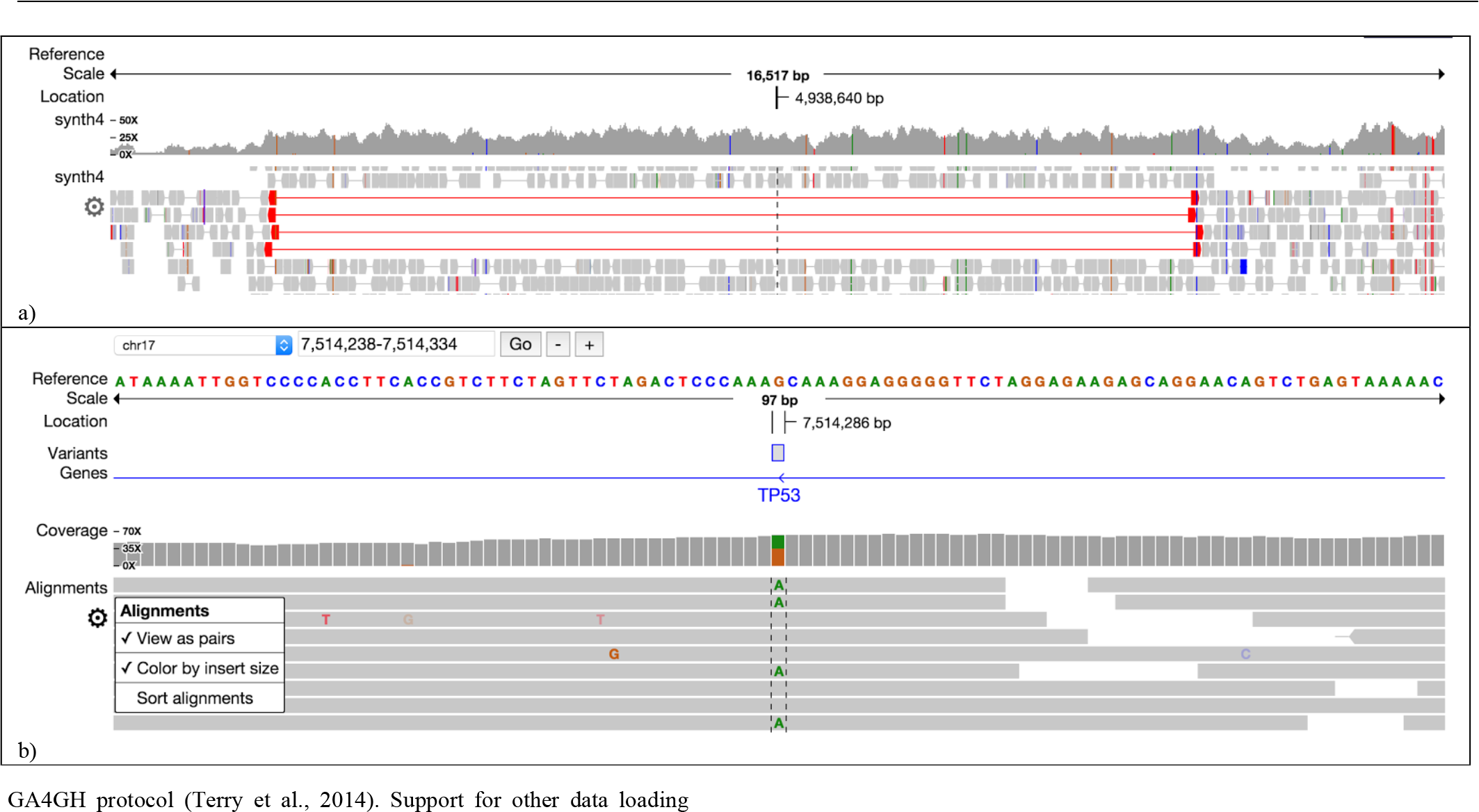
pileup.js. This shows sequencing data at two different zoom levels **(a)** The order and style of the tracks (e.g. reference, variant, gene, coverage and alignment) can be customized. **(b)** The alignment and coverage tracks highlight variants and abnormal reads to draw attention to anomalous regions.

## 3. Features

pileup.js has the standard tracks required for investigating genomic variants (see Figure 1):

1. **Reference track** for visualizing individual base pairs in a reference genome
2. **Gene track** for annotating genomic regions with gene names (introns, exons, coding regions)
3. **Pileup track** for showing (paired or unpaired) aligned sequencing reads
4. **Coverage track** to show the number of reads aligned to each locus, as well as the frequency of variants.
5. **Variant track** for marking regions on the genome containing a called variant.

Additional tracks may be defined by developers using the pileup visualization API. Users can pan and zoom to find and drill down into regions of interest. An options menu allows the view to be configured on a per-track basis, e.g. to view reads individually or as pairs.

The set of tracks and their order can be configured through the JavaScript API. Details of the layout (e.g. track heights and font choices) are designed to be configured via CSS. pileup.js makes use of the UMD (Universal Module Definition) pattern. This allows it to be included in a larger application either via a global variable or as a dependency via a module system such as AMD or CommonJS. It also means that, should they choose to do so, other libraries can depend on just a subset of pileup.js, e.g. its data loading and parsing modules.

pileup.js supports the latest versions of the major browsers at the time of publication: Chrome 42+, Firefox 37+, Safari 9+ and Internet Explorer Edge (12).

We hope that, by providing an easily embedded, modern track viewer, pileup.js will be an enabling tool for the next generation of genomic web applications. External contributions (code and issues) are welcome.

## Acknowledgments

We thank all members of the Hammer Lab for their invaluable input on the manuscript and the tool.

